# An Electrochemistry Study of Cryoelectrolysis in Frozen Physiological Saline

**DOI:** 10.1101/077545

**Authors:** Thomas J. Manuel, Pujita Munnangi, Boris Rubinsky

**Keywords:** Cryoelectrolysis, cryoelectrolytic ablation, cryogenics, cryosurgery, electrolysis, electrolytic ablation, frozen tissue, tissue ablation

## Abstract

This is the first quantitative study on the fundamental physical and electrochemical processes that occur during cryoelectrolysis. Cryoelectrolysis is a new minimally invasive tissue ablation surgical technique that combines the processes of electrolysis and solid/liquid phase transformation (freezing). We measured the pH front propagation and the changes in resistance in a tissue simulant made of physiological saline gel with a pH dye as a function of the sample temperature in the high subzero range above the eutectic. Results demonstrate that effective electrolysis can occur in a high subzero freezing milieu and that the propagation of the pH front is only weakly dependent on temperature. These observations are consistent with a mechanism involving ionic movement through the concentrated saline solution channels between ice crystals at subfreezing temperatures above the eutectic. The Joule heating in these microchannel may cause local microscopic melting, the observed weak dependence of pH front propagation on temperature, and the large changes in resistance with time. In addition, we observed that the pH front propagation from the anode is more rapid than from the cathode. The explanation is the electro-osmotic flow from the cathode to the anode. The findings in this paper may be of fundamental value for designing future cryoelectrolytic ablation surgery protocols.

## I. Introduction

Minimally invasive tissue ablation is an important alternative to surgical resection, often allowing for reduced duration of the surgery and of the subsequent hospitalization, improved access to the surgical site, and optimized treatment of the target tissue [1, 2]. For example, processes such as non-thermal irreversible electroporation [3, 4] and electrolytic ablation [5–7] can be used to ablate cells in a volume of tissue while maintaining important structures in that volume, such as blood vessels, intact. These and many other physical and chemical principles are used as the basis for various minimally invasive tissue ablation techniques, each with their advantages and disadvantages. Recently, our group has become involved in studying novel combinations of these ablation techniques, focusing now on composite technologies combined with electrical components. These combinations include: electrolysis and electroporation, cryosurgery and electroporation, and cryosurgery and electrolysis. This paper pertains specifically to the latter, termed cryoelectrolysis, which is a largely unexplored process.

Cryoelectrolysis is a new minimally invasive tissue ablation surgical technique that employs freezing and electrolysis to ablate undesirable tissues. A recent rise in interest surrounding this technique is marked by its potential to utilize the advantages of both cryosurgery and electrolytic ablation while overcoming their disadvantages. While recent studies on animal tissue have demonstrated the advantages of cryoelectrolytic ablation over cryosurgery ablation and electrolytic ablation used exclusively [8], the fundamental physics of the cryoelectrolytic process have not yet been investigated. This study elucidates a more fundamental understanding of the electro-physical processes which arise with the combination of cryosurgery and electrolysis. First, a brief review on the principles and attributes of cryosurgery and electrolytic ablation when used in minimally invasive surgery is provided below.

### A. Cryosurgery

Cryosurgery is the ablation of undesirable tissues via freezing [9] which typically employs a cryosurgical probe internally cooled by subzero fluid inserted into the undesirable tissue. Freezing propagates from the cryoprobe surface outward ablating the undesirable tissue along the way. The extent of freezing can be monitored in real time via medical imaging techniques which facilitates real time control over the extent of freezing [10–12]. However, it was also found that cells can survive freezing at high subzero freezing temperatures [13, 14]. Therefore, cells can survive on the outer rim of the frozen lesion or around blood vessels. Thus, the extent of freezing seen on medical imaging does not correspond to the extent of cell death. Motivated by the desire to ablate all the cells in the frozen lesion, “combinatorial” cryosurgery is of current interest to researchers in the field [15]. Combinatorial cryosurgery research deals with attempts to enhance frozen cell ablation through additive chemical mechanisms [16–19]. Cryoelectrolysis is one such technique whose goal is to ablate cells surviving freezing in the high subzero freezing range with products of electrolysis. It draws from the technology of electrolytic ablation.

### B. Electrolytic Ablation

Electrolytic ablation is based on the electrochemical phenomena of electrolysis. The physical principles of the process of electrolysis were introduced by Faraday and have been extensively studied since the middle of the 19^th^ century. Electrolysis occurs at the surface of electrodes submerged in an ionic conducting media during the passage of an electric current. New chemical species, generated at the interface of the electrodes as a result of the electric potential driven transfer between electrons and ions or atoms, diffuse away from the electrodes into tissue in a process driven by differences in electro-chemical potential. In physiological solutions, these electrolytic reactions yield changes in pH, resulting in an acidic region near the anode and a basic region near the cathode. The cytotoxic environment developing due to local changes in pH as well as the presence of some of the new chemical species formed during electrolysis cause cell death. Electrolytic ablation, also known as Electro-Chemical Therapy (EChT), employs these product of electrolysis to induce cell ablation [20]. The total amount of electrolytic products generated during electrolysis is proportional to the charge delivered during the process; thus the total charge is used as a quantitative measure of electrolysis. The chemical products at the anode differ from those formed at the cathode resulting in distinct mechanisms of ablation. Electro-osmotic forces drive the migration of water from the anode to the cathode, further magnifying the contrasting physiological effects at the electrode surfaces. Electrolytic ablation requires very low direct currents (10 – 300 mA) and very low voltages (5-25 V) [20]. This is advantageous because it makes the technology simple and safe. However, the ablation procedures are often time consuming, lasting from tens of minutes to hours. The length is related to the slow diffusion of electrochemically produced species in tissue and the concentration dependent rate of cell death inducing electro-chemical reactions. A clinical study on tissue ablation with electrolysis states that— “Currently, a limitation of the technique is that it is time consuming” [6, 7].

### C. Cryoelectrolysis

Cryoelectrolysis was developed with earlier fundamental observations in mind. Research revealed that the freezing of tissue increases the concentration of solutes around cells by removing the water from the solution in the form of ice [21]. Freezing also causes cell membrane lipid phase transition, disrupts the cell membrane lipid bilayer, and causes permeabilization [19]. It occurred to us that the effects of freezing on the cell membrane permeabilization and on increasing the concentration of solutes near the cells should increase exposure of the interior of the cells to the products of electrolysis. These effects would thereby enhance cell death and shorten the time required for electrolytic ablation. The potential effect of these unique features imposed upon the time constraints and efficacy of current ablation methods forms the basis for cryoelectrolytic ablation which manipulates the combination of solid-liquid phase transformation and electrolysis. Experiments in animal tissue have confirmed our hypothesis and have shown that cryoelectrolysis is more effective at cell ablation than either cryosurgery or electrolytic ablation alone [8]. To the best of our knowledge there are no reports on fundamental research focusing on ionic currents in frozen saline. However, knowledge on how ionic currents flow in frozen saline solutions is central to developing optimal cryoelectrolysis protocols. This study reduces this gap in knowledge.

## II. Methods and Materials

A physiological saline based agar was used to simulate tissue. One liter of water was mixed with 9 grams NaCl (Fisher Chemical) and 7 grams of agarose (UltraPure Agarose, Invitrogen). The solution was stirred and heated for 10 minutes and then removed from heat. Two pH indicator dyes were added after five minutes of cooling. For analysis of electrolysis near the anode, methyl red (Sigma-Aldrich®, St. Louis, MO, USA), 1 mL per 100 mL agar solution, was used. Phenolphthalein Solution 0.5 wt. % in Ethanol (Sigma-Aldrich) was used for analysis of electrolysis near the cathode at a concentration of 5 ml per liter agar solution. The color differences seen later between figures 2 and 3 are due to the different dyes present for the alternate configurations. The agar was cast in a 2.5 inch inner diameter, 2.5 inch tall cylindrical plastic vessel. The walls were coated with a 200 *μm* thick copper foil to a height of 1 inch. The height of the gel cast is also 1 inch. A schematic of the setup is shown in Figure 1.

**Fig 1.**
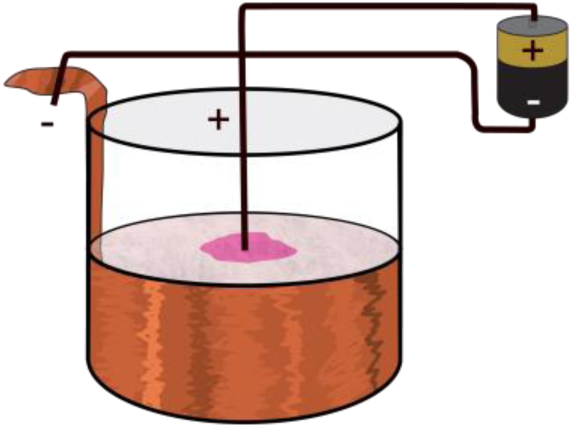
Schematic for anode-centered experiment setup. A 1” deep cylindrical layer of frozen physiological saline was connected to a power supply. The central electrode was a 1mm titanium rod coated with iridium oxide. For cathode-centered experiments, the terminals from the power supply were reversed.

**Fig 2.**
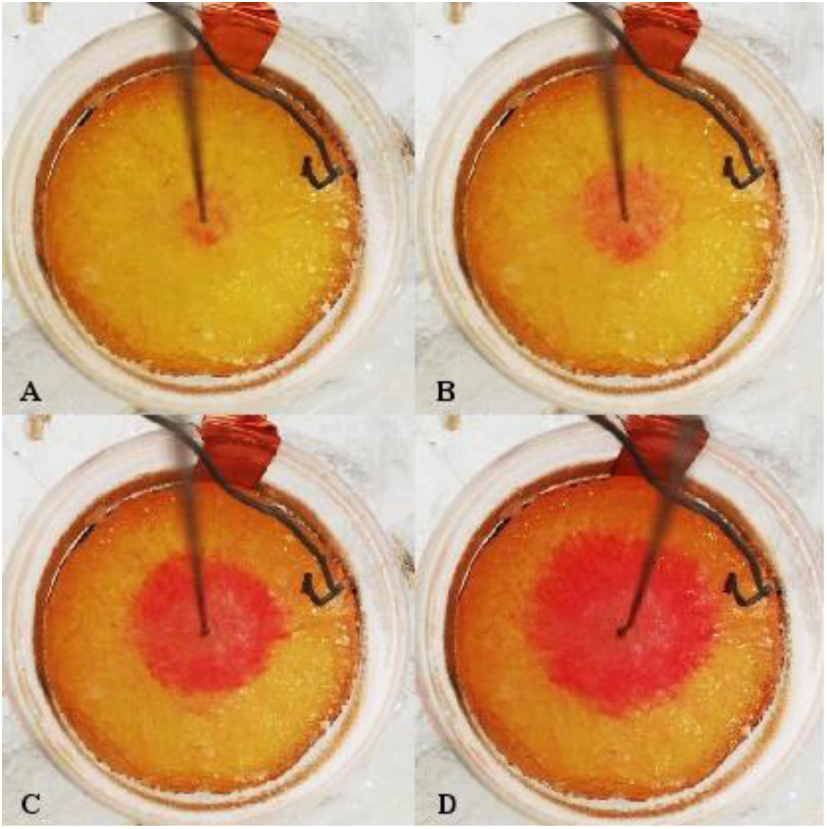
Four photos taken from an anode centered experiment. Note the titanium electrode in the center, the ring of increasing pH propagation, and the 1mm thermistor (curvy wire) used for temperature monitoring.

**Fig 3.**
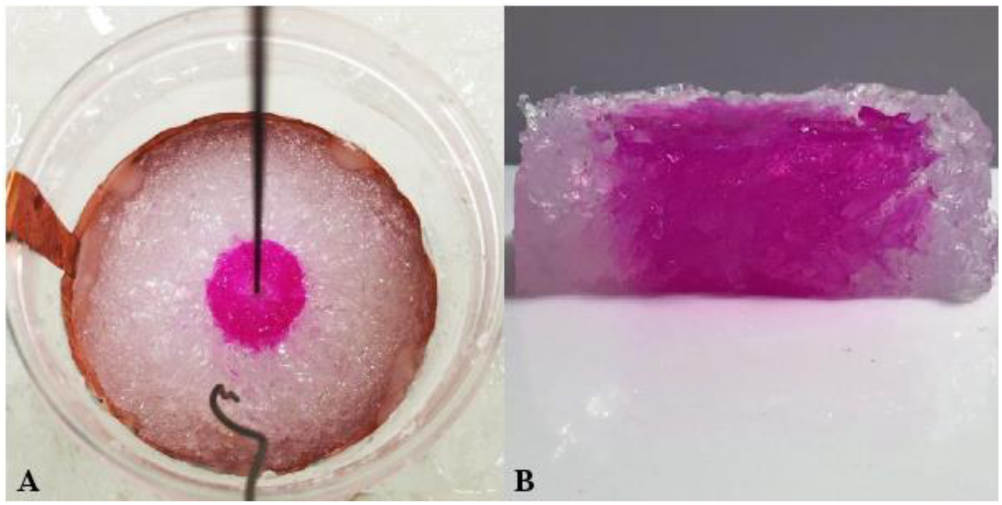
(A) is an image taken from a cathode-centered experiment. (B) is an image showing the cross section of a cathode-centered sample taken after testing displaying the constant propagation throughout the depth of the sample.

To study the process of electrolysis as a function of frozen sample temperature, samples were brought to a desired subzero temperature and held constant for the duration of the experiment. This temperature control was achieved as follows: The jars containing the physiological milieu were sealed with a lid to avoid moisture loss and placed in a freezer at −20 °C overnight to freeze. All samples were initially frozen with identical conditions (free air at a temperature of −20 ̊C). After completing overnight freezing, samples were placed in a constant temperature bath held at the final desired temperature via a mixture of salt and ice. Samples were placed in this constant temperature ice bath in such a way that the top level of the bath was 15 mm above the level of the gel surface. The samples were allowed to achieve thermal equilibrium with the bath prior to the start of the experiment. This ice-bath frozen jar configuration can be seen in Figure 2 (anode centered) as well as Figure 3A (cathode centered). A thermistor was used for temperature monitoring which can be seen in these figures as the curvy wire. The composite temperature measured by the inserted thermocouple remained constant throughout the experiments. Also visible is the central electrode- a 1 mm diameter titanium rod coated with iridium oxide. An Agilent E3631A Triple Output Power Supply was connected to the titanium rod and to the copper tape to power the electrolysis process.

The following experimental procedure was used in all the experiments: Current was initiated once the sample reached thermal equilibrium with the ice bath at the desired temperature. The current and voltage were then continuously recorded. The position of the pH front was also continuously recorded with an iPhone 6 camera iMotion v3.1 (a stop-motion photography application).

Experiments were run at temperatures of – 3 +/−1 °C, −7 +/− °2C, and −12 +/−2 °C. Each temperature group was tested independently at 50 mA and 100 mA. To obtain rates for both anode and cathode pH propagation, the central electrode served as an anode for half of the experiments and as a cathode for the other. Three to four repeats were performed for each configuration with 48 experiments recorded in total. The radius of the pH front, an important datum in our study, was obtained by averaging radius measurements of 14 random points along the perimeter of the propagation per configuration. Measurements were acquired via post-processing image analysis.

## III. Results and Discussion

Figure 2 shows a series of photographs that illustrate several important observations which were typical to all the experiments performed in this study. In these images the propagation in time of an acidic pH front (the red, expanding ring) is clearly visible. Similarly, panel 3A shows the appearance of a basic pH front taken from a cathode centered experiment. Figure 3B shows a cross section from a sample revealing consistency in pH propagation along the central axis of the sample. Figure 4 shows the position of the pH front (radius in mm) in time for the anode-centered and cathode-centered experiments at 100 mA for three different sub-freezing temperature groups. It also displays the total resistance of the samples calculated from voltage and current data. Likewise, figure 5 shows the same values for experiments run at 50 mA.

**Fig 4.**
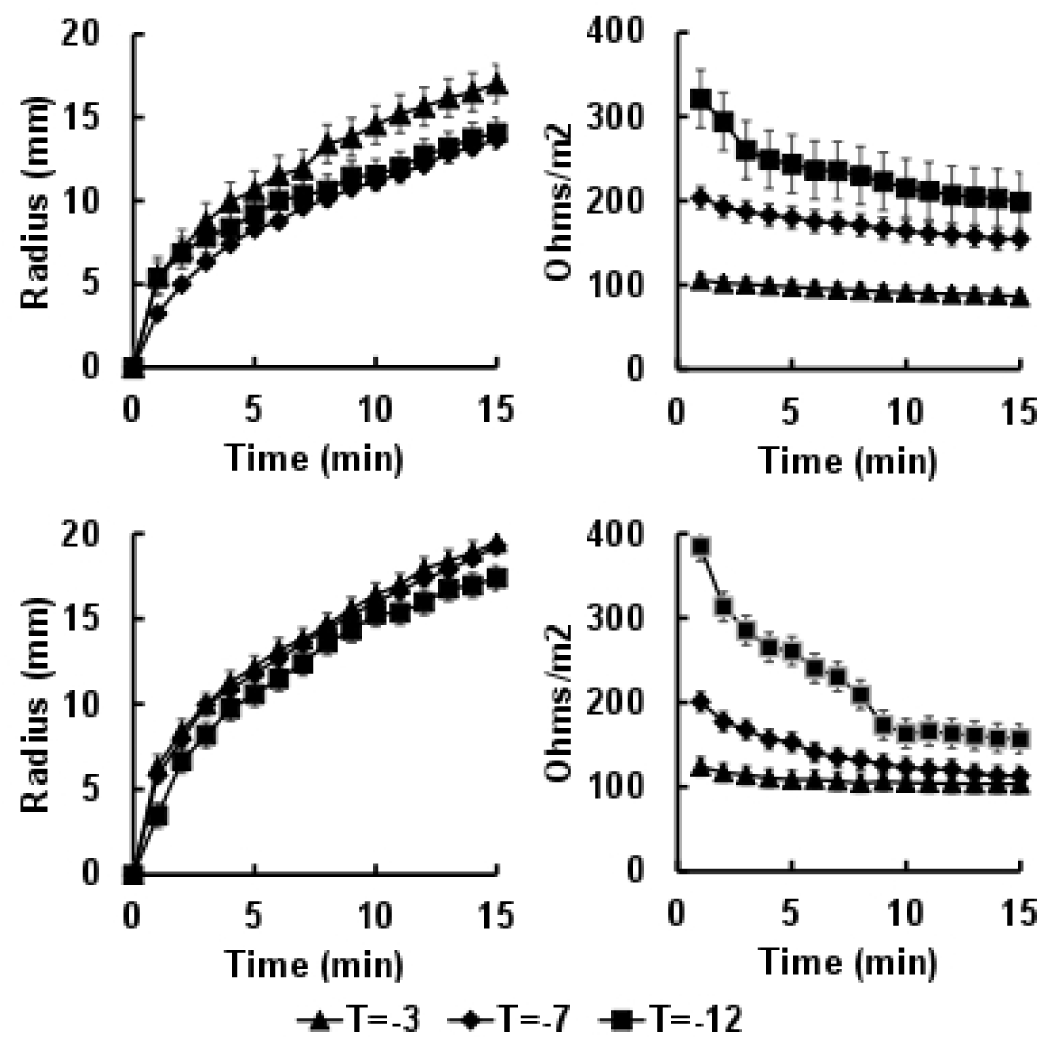
Results from 100 mA experiments at temperatures −3, −7, and −12 degrees Celsius. **Top**: pH front propagation (left) and resistance (right) for cathode-centered experiments. **Bottom**: pH front propagation (left) and resistance (right) for anode-centered experiments.

**Fig 5.**
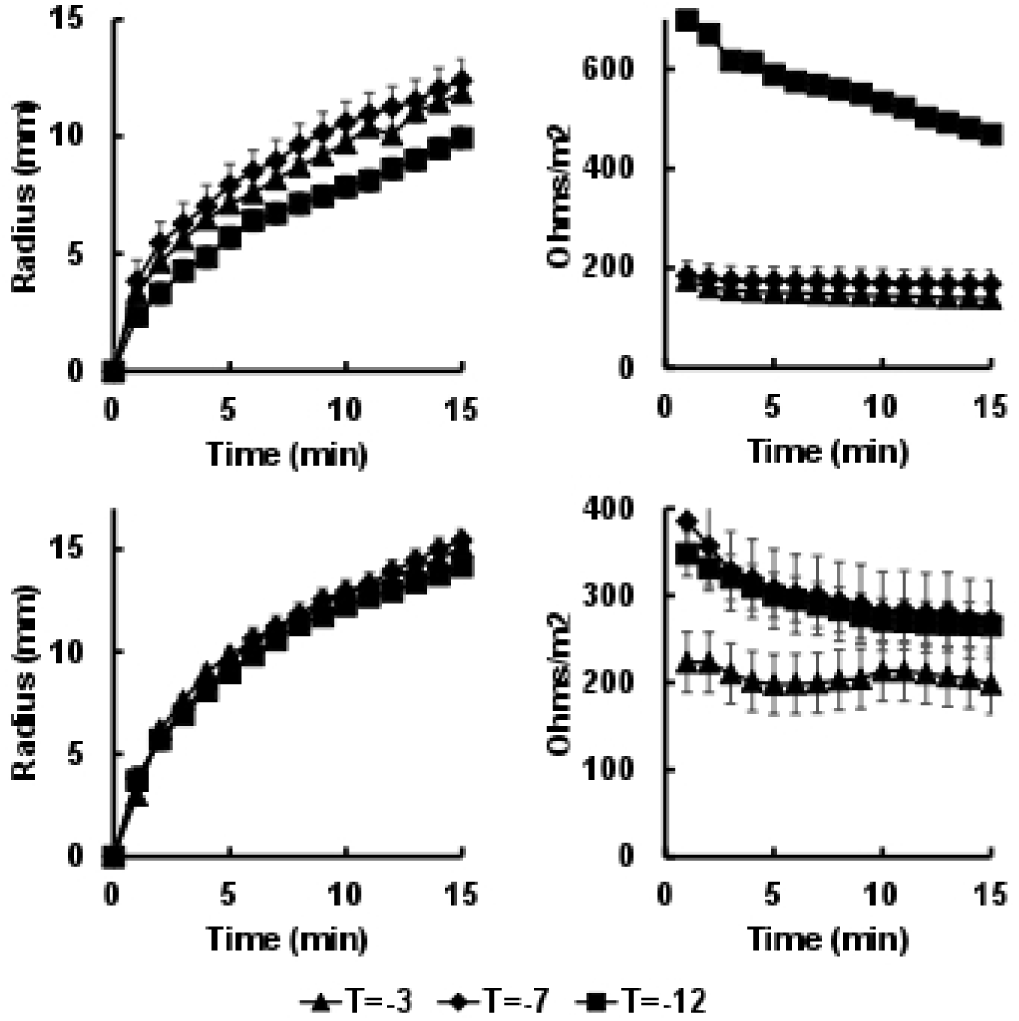
Results from 50 mA experiments at temperatures −3, −7, and −12 degrees Celsius. **Top**: pH front propagation (left) and resistance (right) for cathode-centered experiments. **Bottom**: pH front propagation (left) and resistance (right) for anode-centered experiments.

Figures 4 and 5 provide novel proof that electrolysis can occur, currents can flow, and that pH fronts will propagate through a frozen physiological saline milieu. They also supply quantitative data on the rate at which the process happens for currents of 50 and 100 mA. From the results, it is clear that the propagation rates are mostly proportional to current supply and only minimally dependent upon the temperature of the tissue simulant. This information is of fundamental value for the feasibility of cryoelectrolytic devices, and provides critical values to form a basis for future cryoelectrolytic protocols.

Further analysis was taken in attempt to reveal the mechanism of electrolysis through the frozen gel. Observe that for warmer subzero temperatures the total resistances are relatively constant throughout the process. However, for cooler subzero freezing temperatures, the resistance decreases with the time of electrolytic activation. This observation is consistent with for both Figures 4 and 5. The key to understanding the mechanism responsible for electrolysis in the frozen tissue simulant lies in the fact that the resistance remained constant in time for warmer subzero temperatures but decreased in time for cooler subzero temperatures.

Figure 6 in conjunction with Figure 3B provides an explanation for this observation. Figure 6 is a freeze-dried structure of a frozen agarose gel (from [22] with permission). The important observation is that the frozen agar structure is rugged with finger-like salt/agar structures that formed between the ice crystal during freezing. These finger-like structures become colored in our experiments following the pH-sensitive dye color activation. This can be seen in Figure 3B which shows a cross section of a cathode-centered sample taken after an experiment. We hypothesize that the mechanism responsible for electrolysis in a high subzero frozen media is associated with the specifics of frozen interface propagation in solutions and tissues. The crystallographic structure of ice is tight and cannot incorporate solutes. Fundamental principles related to the instability of the ice interface due to constitutional supercooling demand that during the freezing of a solution finger-like ice crystals form, the salt is rejected along these ice crystals [23], and high concentration salt solution layers manifest between the ice crystals. This phenomenon occurs during the freezing of any aqueous medium, in solutions[24], in gels[22] and in tissues [21]. While the electrical conductivity of ice is essentially zero, electrical currents can flow through these high concentration brine channels until the temperature reaches the eutectic − 21.1 °C. Figures 2 and 3 show that, indeed, current can flow through the physiological milieu at high subzero temperatures. Unavoidably, this flow of ionic current initiates electrolysis, driving the pH front's advancement through the frozen tissue simulant.

**Fig 6.**
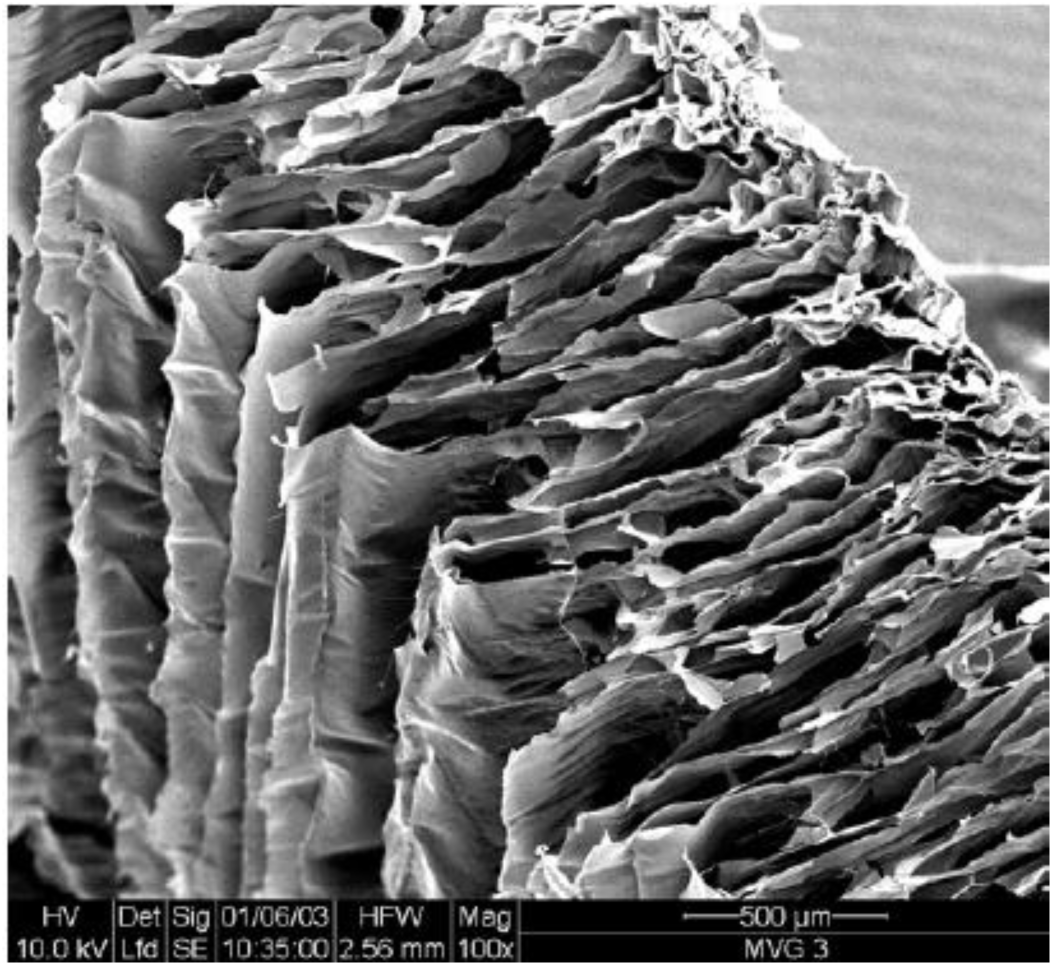
SEM image of freeze-dried agar. Visible structures are salt/agar solids that formed due to the freezing of the agar.

The observed decrease in resistance with time may then be explained by local Joule heating effects within the microscopic brine channels. The flow of current through these brine channels most likely elevates the local temperature, causes local melting, and results in either expansion or collapse of the brine channels. Any Joule heating must be a local microscopic effect, seeing that the bulk temperature measured via the thermistor remained constant throughout the experiments. This may be of fundamental as well as practical value in designing future cryoelectrolysis protocols as it indicates that the cryoelectrolytic process will be essentially independent of the temperature of the frozen lesion as long as it is warmer than the eutectic.

Figure 7 illustrates a final important aspect of the electrolytic process in a frozen tissue simulant. Rather than compare pH propagation rates for different temperatures, Figure 7 plots propagation rates from the anode versus those from the cathode. Standard error bars are invisible due to their small values (<0.6 mm). Results indicate that for the same current, the pH front propagates faster from the cathode than the anode. The propagation of the pH front is driven by diffusion but also by electro-osmosis which occurs from the anode to the cathode- a distinction responsible for the difference in the pH front propagation between the anode and cathode. While the difference between the process of electrolysis at the anode and cathode is well understood [1, 2], Figure 7 shows that these factors will also need to be considered in designing optimal cryoelectrolysis protocols.

**Fig 7.**
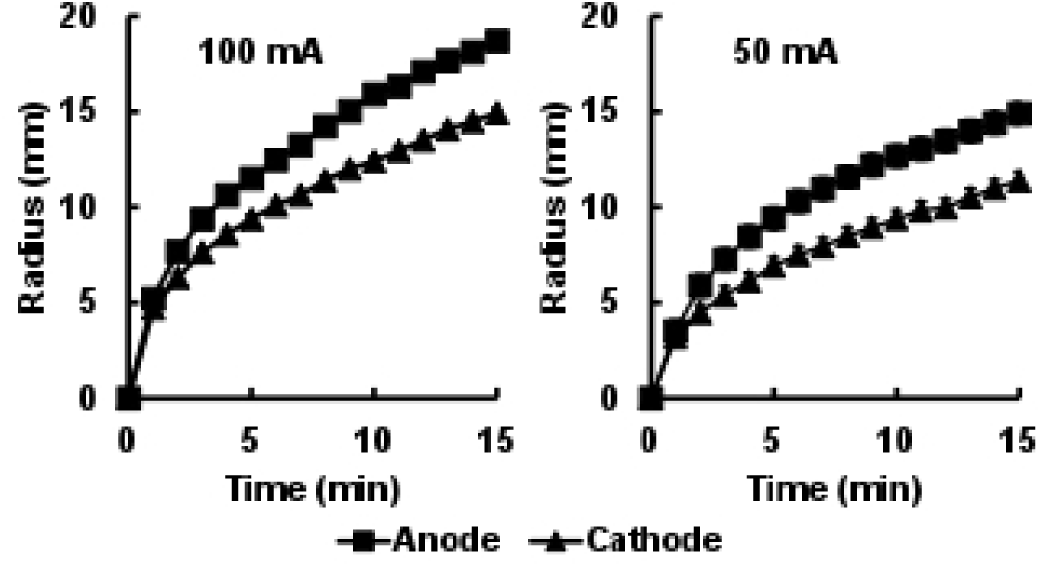
pH propagation comparing anode rate to cathode rate for 100 mA (left) and 50 mA (right) of all temperature ranges (T=−3, −7 and −12 °C).

## IV. Conclusion

In summary, our study demonstrates and quantifies three important physical phenomena related to cryoelectrolysis:

a. *Electrolysis can occur through a frozen milieu at both the anode and the cathode.*
b. *The rate of the resulting pH front propagation is only weekly affected by the temperature of the frozen milieu.*
c. *The propagation of the pH front is strongly affected by the electro-osmotic flow from the anode to the cathode and will need to be considered in designing cryoelectrolysis protocol.*

To the best of our knowledge, this is the first time that a systematic study of the effects of freezing temperature on electrolysis through ice was performed and reported. The findings support an optimistic future for cryoelectrolysis, quantitatively demonstrating the effectiveness of combining cryogenics and electrolysis, and demonstrating a minimal process rate dependence on the low subzero temperature ranges which will be required.

